## Supplementary Information for "Organization of fast and slow chromatin revealed by single-nucleosome dynamics"

**This PDF file includes:**

Supplementary text
Figs. S1 to S9
Caption for Movie S1
References for SI reference citations

**Other supplementary materials for this manuscript include the following:**

Movie S1

### Supporting Information Text

**Single-nucleosome trajectories.** In the microscopic observations of Nozaki et al. (? ), photoactivatable (PA)-mCherry (? ) was fused with histone H2B and expressed in HeLa cells. The oblique illumination method (? ? ) was used to illuminate a thin layer ($\sim$ 200–250 nm thickness) in a single nucleus. A relatively small number ($\sim$ 100 molecules/frame) of H2B-PA-mCherry molecules were continuously and stochastically activated without UV laser stimulation (see Movie S1). The imaging frame was taken every 50 ms and $\sim$ 1000 frames ($\sim$ 50 s in total) were obtained. In a photoactivated localization microscopy (PALM) image constructed from these frames, a total of $\sim 20,000$ single nucleosomes were detected as fluorescent dots, amounting to $\sim 2\%$ of the total number of nucleosomes in the layer. $\sim$ 100 dots newly appeared in every frame and disappeared after $\sim$ 0.05–1 s (1–20 frames). An example image of a frame and the PALM image are shown in Fig. S1 and Movie S1.

The fluorescence intensity of the observed dots disappeared with a single-step photobleaching in a quantized manner; the fluorescent intensity of dots dropped from approximately 70 (in arbitrary units) to 0 within 50 ms, suggesting that each dot represents a single H2B-PA-mCherry molecule in a single nucleosome (for details, see Figure S1B in Ref. 1). With the time frame resolution being $t = 0.05$ s, shown in Fig. S2A is $G_s(r, t = 0.05\,\mathrm{s})$, which results in an average displacement of each nucleosome to be $\sim$ 50 nm in this time frame, with $> 95$ % of the particles having displacements $< 100$ nm (shown in Fig. S2B). Shown in Fig. S2C is the pair correlation function for the nucleosomes (without distinguishing between fast and slow), which indicates infrequency of finding particles within 200 nm. This shows that the particles are spatially well resolved and can be faithfully labelled without any confusion from frame to frame. See Movie S1 showing movements of the fluorescent PA-mCherry dots.

Long-time laser illumination may damage DNA or denature proteins. Indeed, the average MSD in the last 200 frames among $\sim$ 1000 frames observed is smaller than that in the first $\sim$ 500 frames, showing the increase of slow nucleosomes through the laser illumination. Similarly, by using the laser with the stronger intensity, the MSD is smaller with the larger population of the slow nucleosomes. In order to minimize the effects of DNA damage, we used the first 500 frames (25 s total) for the analyses in the present study. From the observed $\sim 20,000$ nucleosome trajectories, we selected trajectories which last longer than 0.5 s, resulting in $\sim$ 1–2 nucleosomes in one frame and 300–600 trajectories during 25 s. For calculating pair correlations, we used pairs that appeared simultaneously in the same frame, which resulted in the sample size $\sim$ 1 pair/frame and 400–500 pairs in 25 s.

**Effects of 2D projection.** The observed trajectories are those projected on the 2D imaging plane. The 2D projected MSD of Fig. 1B is related to the MSD of the 3D movement $M^{\mathrm{3D}}$ as $M = \gamma M^{\mathrm{3D}}$, where the factor $\gamma \sim 2/3$ represents the effect of reducing the spatial dimension from 3 to 2 for isotropic dynamics. The movement of the nucleosomes at the nuclear periphery can be anisotropic to show $1/2 \lesssim \gamma < 2/3$. However, the population of nucleosomes near the periphery is small and gives only a minor contribution to $P(M, t)$. Therefore, the peak of slow nucleosomes in $P(M, t)$ does not result from the projection but from the intrinsic mechanism that is distinct from the movement of fast nucleosomes. The further discussion on the effects of the projection is given with a ring polymer model, whose results are shown in Fig. S3.

**A ring polymer model.** A ring polymer chain was used for calculating the results shown in Fig. 6 of the main text and Fig. S3. The chain was represented by a bead-and-spring model with 300 beads. The chain dynamics was simulated with the interaction potential between $i$ and $j$th beads,

$$V(r_{ij}) = \epsilon_0 (r_0/r_{ij})^{12} - \epsilon_{ij} (r_0/r_{ij})^6. \qquad \text{[S1]}$$

We used an $r_{\mathrm{cut}} = 3.5 r_0$; for $r_{ij} > r_{\mathrm{cut}}$, $V(r_{ij}) = 0$. The finite extensive nonlinear elastic (FENE) potential was assumed for the spring between $i$ and $i+1$th beads, which provides a force,

$$F(r_{ii+1}) = k(r_0 - r_{ij})/(1 - u_b^2). \qquad \text{[S2]}$$

Here, $u_b = (r_0 - r_{ij})/r_0$ and $k = 1$. We used the overdamped Langevin dynamics to simulate the system with the friction coefficient $\gamma = 1.0$ and temperature $k_{\mathrm{B}}T = 1.0$. We set $r_0 = 1.0$ and $\epsilon_0 = 1.0$. The integration time step was $\Delta t = 0.001$. We let the system come to equilibrium after which we collected statistics of MSD at $t_m = 5000\Delta t$.

***A chain confined in a sphere: examination of the projection effects.*** The ring chain described above was used with $\epsilon_{ij} = 1$ for all $i$ and $j$. The chain was confined in a sphere of radius $9 r_0$ with a repulsive potential

$$V_{\mathrm{wall}}(r) = \epsilon_w \left((r_s - r)/r_s\right)^5, \qquad \text{[S3]}$$

where $r_s = 2r_0$ and $r$ is the distance of the bead to the wall of the sphere. $\epsilon_w = 10.0 k_{\mathrm{B}}T$. Shown in Fig. S3A is a snap shot of the simulated system and its projection onto the 2D plane. Shown in Fig. S3B is the histogram of the MSD at $t_m$ calculated directly from the 3D trajectories of individual beads, and the smooth curve is $P(M) = P(M, t_m)$ calculated by using the RL algorithm in which the van Hove correlation function calculated from the 3D coordinates of beads was used. The histogram and the distribution $P(M)$ of MSD were also calculated for the 2D projected dynamics as shown in Fig. S3C. We see both in 2D and 3D, the RL algorithm works well to quantitatively reproduce the MSD distribution directly calculated from the trajectories. The MSD in the 2D space is about 2/3 of the MSD in the 3D space. The effects of anistropy are not significant in the results of the 2D projection.

***A chain used for calculating the results in Figure 6.*** Region I and Region II were defined in the ring described above; Region I is from $i = 1$ to 100 and Region II is from 101 to 300. This ring chain model was used with $\epsilon_{ij} = \epsilon_{\rm I}$ when $i$ and $j \in$ Region I, $\epsilon_{ij} = \epsilon_{\rm II}$ when $i$ and $j \in$ Region II, and $\epsilon_{ij} = 0$ when $i$ and $j$ belong to different regions in the ring. A strong force $\epsilon_{ij} = 100k_{\rm B}T$ was assumed between the beads $i = 1$ to 101 to define the region boundaries. The ring was placed in the open space without confinement and the origin of the coordinate (reference point) was fixed on the $i = 50$th bead in Region I or 150th bead in Region II.

**Mapping fast and slow nucleosomes.** In the main text, the nucleosomes are characterized as fast and slow based on the single nucleosome MSD distribution derived with the RL algorithm. Here, we check the consistency of this characterization by using a polymer model; we compare the results from the RL algorithm and the results directly calculated from the trajectories of individual beads in a linear chain polymer.

The polymer chain has 100 beads, which are connected to the neighbors via springs with a spring constant $k = 3.0$. Interactions between beads, $V(r_{ij})$, are assumed as

$$V(r_{ij}) = \epsilon_0 (r_0/r_{ij})^{12} - \epsilon_{ij}(r_0/r_{ij})^6, \qquad \text{[S4]}$$

We used $r_0 = 1$ and $\epsilon_0 = 1$. The beads $i > N_c = 30$ are in the extended part and beads $i \le N_c$ are in the compact part. We assumed $\epsilon_{ij} = 0$ when $i$ or $j$ is larger than $N_c$, and examined two cases; (a) $\epsilon_{ij} = 10$ for $i, j \le N_c$ and (b) $\epsilon_{ij} = 3$ for $i, j \le N_c$. We used the overdamped Langevin dynamics to simulate the system with the friction coefficient $\gamma = 1.0$ and temperature $k_{\rm B}T = 1.0$. The integration time step was $\Delta t = 0.001$. The system was first brought to equilibrium and the MSD distribution of single bead trajectories was calculated at $t_m = 5000\Delta t$. The chain was in the open space without confinement and the MSD was calculated by using the coordinate whose origin is fixed on the center of mass of the chain.

Shown in Fig. S4A is a snap shot of the simulated polymer having the compact region (red) and the extended region (green). In Figs. S4B and S4C, the MSD distribution $P(M) = P(M, t_m)$ calculated with the RL algorithm is compared with the histogram of MSD directly calculated from trajectories of individual beads. The bimodal $P(M)$ is split as fast and slow at $M^* = 0.25r_0^2$. We then calculate the split $G_s^{f,s}(r,t)$ for the fast and slow beads as discussed in the main section using the RL algorithm. In Figs. S4D and S4E, the split $G_s^{f,s}(r,t)$ calculated with the RL algorithm is compared with $G_s^c(r,t)$ and $G_s^e(r,t)$ calculated from trajectories of beads in the compact and extended parts of the polymer, respectively. The $G_s^{c,e}(r,t)$ directly calculated from the simulated trajectories match up with the $G_s^{f,s}(r,t)$ calculated with the RL algorithm.

**Robustness of the RL evaluation.** In the single-nucleosome analyses in the main text, we examined $P(M, t_m)$ of the mean square displacement, MSD, with $t_m = 0.5$ s by using trajectories of various length $t_L \sim 0.8$–3.0 s for individual nucleosomes in an example cell. For a precise estimation of the MSD distribution, the mean-value statistics along a long enough trajectory is required. With short trajectories of $t_L/t_m \sim 1.6$–6.0, the mean-value calculation includes statistical errors, which makes distribution of $M$s noisy. However, the RL algorithm filters out this noise. Here, we examine the efficiency of the RL method to obtain the smooth $P(M, t_m)$ from short trajectories by using the same polymer model as used in Fig. S4. We adopt $\epsilon_{ij} = 1.5$ for $i, j \le N_c$ and set $t_m = 5000\Delta t$.

Starting from the polymer model of Fig. S4, long trajectories of length $6t_m$ were generated for the 300 particles in the model. Then, short time intervals $t_L$ were chosen from these long trajectories to mimic those of the experiment; short time distributed intervals for 300 particles were selected to have the same $t_L/t_m$ of the observed 300 randomly chosen nucleosome trajectories in an example cell. As shown in Fig. S5A for the short trajectories, the directly calculated histograms are too noisy to show the bimodal feature, unlike the $P(M)$ (black) from the RL algorithm. In Fig. S5B, we show that the distribution of $M$s directly calculated from the long single-particle trajectories and $P(M)$ calculated with the RL method coincide with each other showing the bimodal feature; the RL-algorithm is sensitive to this bimodal feature even in the short trajectories, and this feature is preserved in the long trajectories. The position $M^*$ of the minimum of $P(M)$ from the RL algorithm, which is used to distinguish the fast and the slow particles, is the same for both short distributed and long trajectories. Further, the area under the curves of the fast and slow peaks in $P(M)$, which indicates the number of fast and slow particles is almost the same ($\sim 3\%$ difference in the ratio of fast and slow particles) for the short and long trajectories. Shown in Fig. S5C is a comparison of $P(M)$ from the long (black) and short (red) trajectories. Thus, this model calculation demonstrates that the RL method is robust against errors in the parameter of interest induced by the short sampled trajectories, and the method is efficient enough for further analyses.

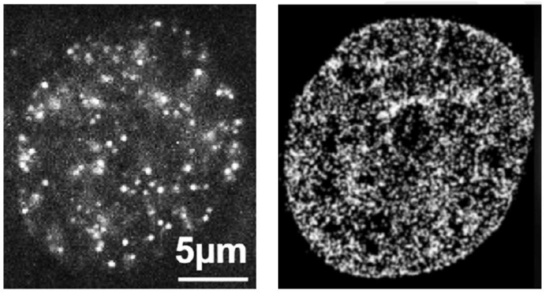


**Fig. S1.** Super-resolution images of the nucleus of a live HeLa cell. (Left) An example frame image of single nucleosomes (H2B-PA-mCherry). (Right) An example PALM image constructed from 1000 frames. Reprinted from Figure 1 of Nozaki et al., (2017) $Molecular\ Cell$ 67: 282–293 with permission. .

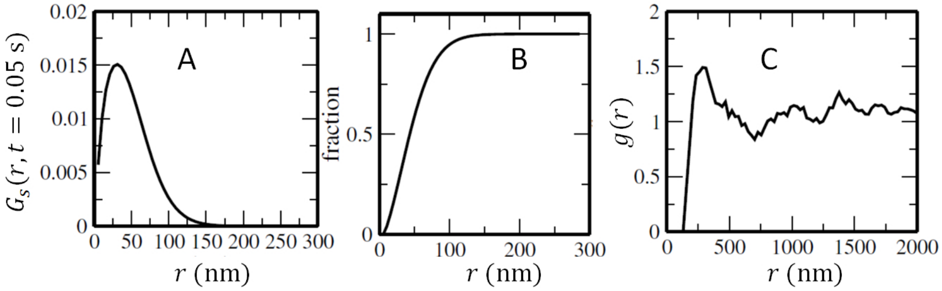


**Fig. S2.** Nucleosome pair is well separated from each other in the observed cells. (A) The van Hove correlation $G_s(r, t = 0.05\,\mathrm{s})$, (B) cumulative function of $G_s(r, t = 0.05\,\mathrm{s})$, and (C) nucleosome pair correlation function $g(r)$. $g(r)$ was calculated by sampling pairs of nucelosomes irrespective of fast or slow.

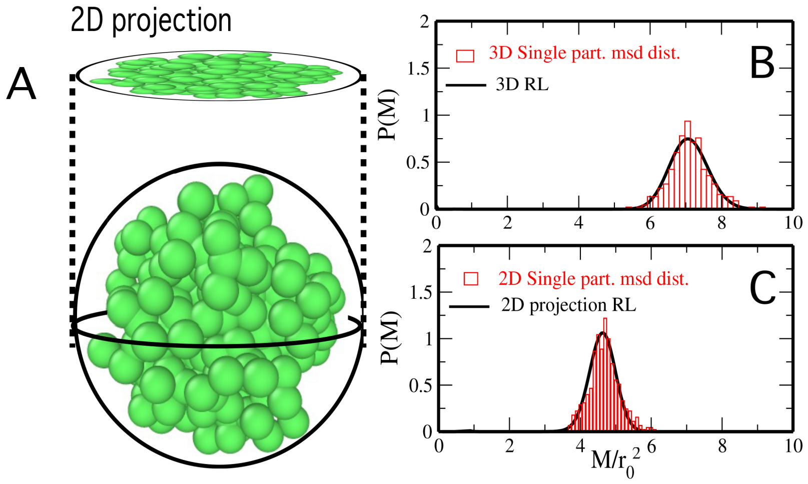


**Fig. S3.** Calculation of the distribution of mean square displacement (MSD), $M$, of beads in a ring polymer chain. (A) A ring polymer is confined in a sphere. Beads of a snapshot structure of the polymer are shown with green spheres. The MSD distribution is calculated with the 3D coordinates and the coordinates in the projected 2D space. (B) Comparison of the distribution $P(M) = P(M, t_0)$ derived with the Richardson-Lucy (RL) algorithm used in the 3D space and the histogram of the MSD distribution directly calculated from the trajectories of individual beads in the 3D space. (C) Comparison of the distribution $P(M) = P(M, t_0)$ derived with the RL algorithm used in the 2D space and the histogram of the MSD distribution directly calculated from the trajectories of individual beads in the 2D space. The distribution calculated with the RL algorithm is in good agreement with the histogram of MSD directly calculated from the trajectories both in the 2D and 3D spaces, showing that the RL algorithm works well for evaluating the distribution of MSD. The MSD calculated in the projected 2D space is about $2/3$ of the MSD calculated in the 3D space, and the distribution of MSD in the 2D space is close to the one in the 3D space when $M$ is scaled with the factor $2/3$.

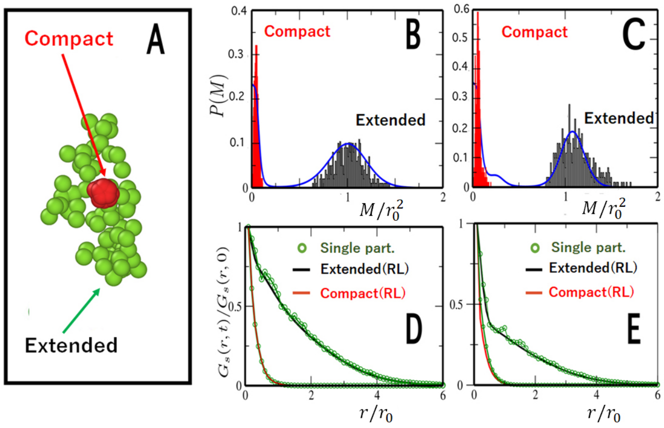


**Fig. S4.** Simulations with a linear polymer model performed to check the consistency in calculating $P(M,t)$ and $G_s^{f,s}(r,t)$. (A) A snapshot of the polymer bead configuration with extended ($i > N_c = 30$, green) and compact ($i \le N_c$, red) parts. (B) The MSD histogram directly calculated from the trajectories of beads in the compact (red) and extended (gray) parts, respectively, and $P(M) = P(M, t_m)$ derived from the RL algorithm in the case $\epsilon_{ij} = 10$ for $i, j \le N_c$. (C) Same as in B with $\epsilon_{ij} = 3$. (D) $G_s^s(r,t)$ (red) and $G_s^f(r,t)$ (black) derived from the RL algorithm are shown by smooth curves. The $G_s^c(r,t)$ and $G_s^e(r,t)$ calculated directly from the trajectories of beads in the compact and extended parts, respectively, are shown by green circles. $\epsilon_{ij} = 10$ for $i, j \le N_c$. (E) Same as in D with $\epsilon_{ij} = 3$.

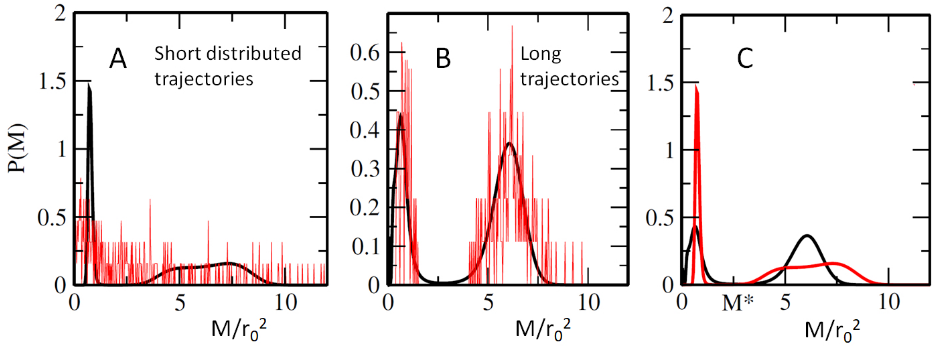


**Fig. S5.** The MSD distribution $P(M) = P(M, t_m)$ calculated from trajectories of various length. (A) The distribution histogram of $M$s directly calculated from the single-particle trajectories whose length $t_L$ distribution was chosen to make the ratio $t_L/t_m$ same as that in the experimental statistics (red) is compared with $P(M)$ calculated with the RL method, $P^{\mathrm{RL}}_{\mathrm{Short}}(M)$ (black). (B) The distribution histogram directly calculated from the single-particle trajectories of length $t_L/t_m = 6$ (red), is compared with $P(M)$ calculated with the RL method, $P^{\mathrm{RL}}_{\mathrm{Long}}(M)$ (black). (C) $P^{\mathrm{RL}}_{\mathrm{Short}}(M)$ (red) and $P^{\mathrm{RL}}_{\mathrm{Long}}(M)$ (black) are superposed. $P^{\mathrm{RL}}_{\mathrm{Short}}(M)$ and $P^{\mathrm{RL}}_{\mathrm{Long}}(M)$ have the minimum at the same $M = M^*$, and the ratio (area under the fast peak)/(area under the slow peak) is 1.85 in $P^{\mathrm{RL}}_{\mathrm{Short}}(M)$ and 1.92 in $P^{\mathrm{RL}}_{\mathrm{Long}}(M)$. The same polymer model as in Fig. S4 was used with $\epsilon_{ij} = 1.5$ for $i, j \le N_c$.

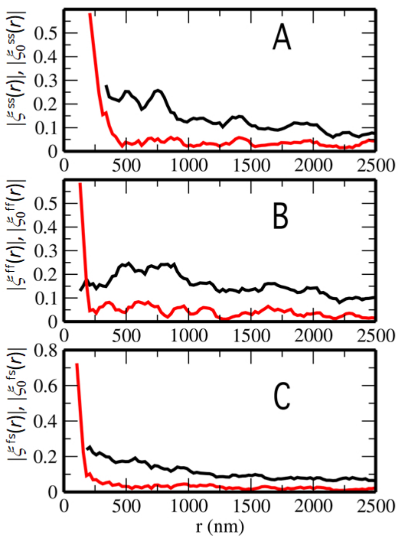


**Fig. S6.** Pair correlation functions of direction of movement of nucleosomes separated at distance $r$, $\xi^{ab}(r) = \left[\xi^{ab}_{vv}(r)\right]_{\rm cell} / \left[\xi^{ab}_{rr}(r)\right]_{\rm cell}$ (red) and $\xi^{ab}_0(r) = \left[\xi^{ab}_{vv}(r)/\xi^{ab}_{rr}(r)\right]_{\rm cell}$ (black). (A) $|\xi^{ss}(r)|$ and $|\xi^{ss}_0(r)|$, (B) $|\xi^{ff}(r)|$ and $|\xi^{ff}_0(r)|$, and (C) $|\xi^{fs}(r)|$ and $|\xi^{fs}_0(r)|$. $|\xi^{ab}(r)|$ has a large value only in the range of $r < R^{ab}_c \approx 190$–300 nm, while $|\xi^{ab}_0(r)|$ shows a micrometer scale correlation.

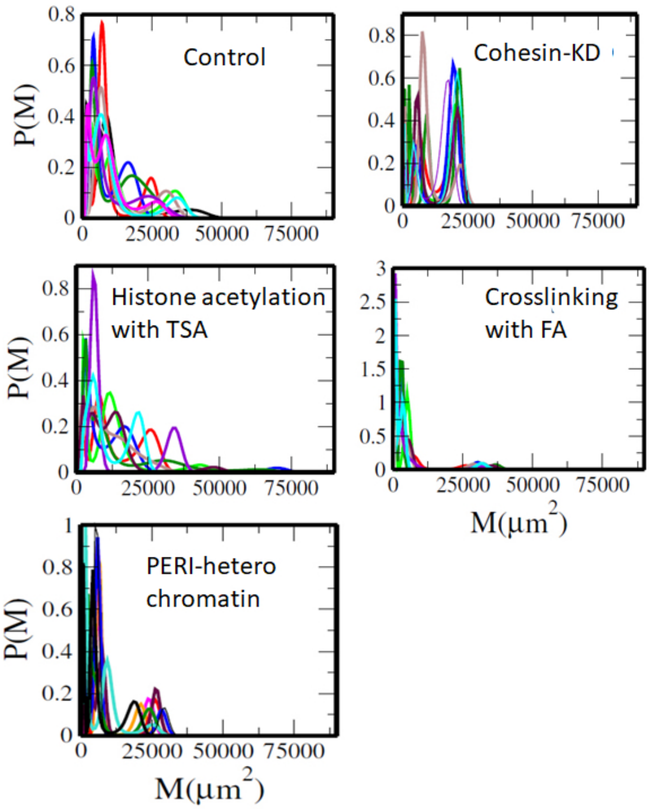


**Fig. S7.** Effects of perturbations and effects of focusing on heterochromatin on the distributions of MSD, $P(M, 0.5\ {\rm s})$, of 10 cells. 10 lines of $P(M, 0.5\ {\rm s})$ are superposed in each case.

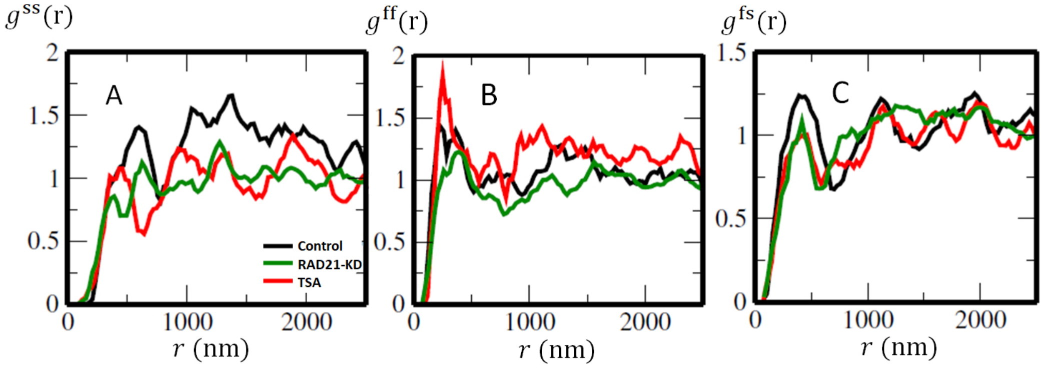


**Fig. S8.** Effects of cohesin-KD (RAD21 knocking down) and histone acetylation with TSA on the radial distribution $g^{ab}(r)$ of nucleosomes. (A) $g^{ss}(r)$, (B) $g^{ff}(r)$ and (C) $g^{fs}(r)$, where $g^{ab}(r)$ are drawn for control (black), cohesin-KD (green), and TSA added (red) cases.

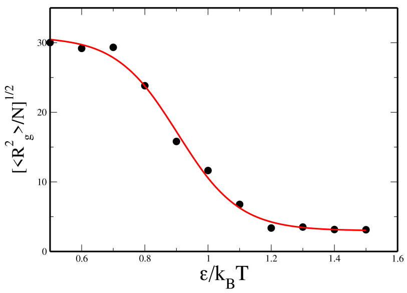


**Fig. S9.** Radius of gyration, $R_g$, of the simulated ring polymer chain having $N = 300$ beads is shown as a function of the strength of attractive interactions, $\varepsilon$. The same ring polymer model as used in Fig. 6 of the main text was used by setting the chain bundling energy mimicking cohesin binding zero. The red curve is a guide for eyes.

**Movie S1. Raw video of single nucleosomes in the living HeLa cell. From Nozaki et al., (2017)** ***Molecular Cell*** **67: 282–293.**
